# Increased volatile thiol release during beer fermentation using constructed interspecies yeast hybrids

**DOI:** 10.1101/2022.06.12.495761

**Authors:** Kristoffer Krogerus, Nils Rettberg, Brian Gibson

## Abstract

Interspecies hybridization has been shown to be a powerful tool for developing and improving brewing yeast in a number of industry-relevant respects. Thanks to the popularity of heavily hopped ‘India Pale Ale’-style beers, there is an increased demand from brewers for strains that can boost hop aroma. Here, we explored whether hybridization could be used to construct strains with an enhanced ability to release hop-derived flavours through β-lyase activity, which releases desirable volatile thiols. Wild *Saccharomyces* strains were shown to possess high β-lyase activity compared to brewing strains, however, they also produced phenolic off-flavours (POF) and showed poor attenuation. To overcome these limitations, interspecies hybrids were constructed by crossing pairs of one of three brewing and one of three wild *Saccharomyces* strains (*S. uvarum* and *S. eubayanus*). Hybrids were screened for fermentation ability and β-lyase activity, and selected hybrids showed improved fermentation and formation of both volatile thiols (4MMP, 3MH and 3MH acetate) and aroma-active esters compared to the parent strains. Undesirable traits (e.g. POF) could be removed from the hybrid by sporulation. To conclude, it was possible to boost the release of desirable hop-derived thiols in brewing yeast by hybridization with wild yeast. This allows production of beer with boosted hop aroma with less hops (thus improving sustainability issues).

## Introduction

The interest toward heavily hopped ale beer and beer with novel and diverse flavours has increased considerably the past years [1–3]. For instance, hop production and consumption within the brewery in the USA has grown markedly in the past decade [4, 5]. Interest towards the use of atypical yeast strains as a strategy for diversifying flavour has also grown [6–8]. Indeed, beer flavour varies considerably depending on the brewing yeast strain employed for fermentation. However, the different flavour profiles tend to vary in a quantitative rather than qualitative way, with brewing yeast producing similar flavour compounds (e.g. higher alcohols, acetate esters, ethyl esters), albeit in different quantities. To diversify and introduce new flavours, as well as meet the demand for enhanced hop flavour, brewers have begun exploiting the ability of yeast to enzymatically alter hop-derived compounds in the beer [9, 10]. This process is often referred to as ‘biotransformation’, and the term encompasses many different types of reactions. Among the key biotransformation reactions are those catalysed by β-lyase enzymes. These enzymes can liberate volatile thiols from their conjugated (glutathionylated or cysteinylated) and therefore non-active forms [11–13]. Volatile thiols, such as 4-mercapto-4-methyl-2-pentanone (4MMP), 3-mercapto-1-hexanol (3MH), and 3-mercaptohexylacetate (3MHA), have been shown to have a vital role in modern hop aroma. These compounds contribute desirable tones of black currant, grapefruit, and passion fruit, and have extremely low flavour thresholds, often in the ng/L range [14].

While some hop varieties may directly contribute considerable amounts of free thiols, they, together with malt, have also been shown to contain high concentrations of conjugated 3MH and 4MMP precursors [11]. Indeed, the vast majority of all thiols in hops, are present in cysteine-or glutathione-conjugated form and are therefore odourless [11]. Hence, there exists a large potential pool of flavour that can be unmasked through the use of yeast expressing the β-lyase activity. This would allow for the production of beer with detectable thiol levels using hop varieties that are otherwise low in free thiols (e.g. Perle or Cascade; Roland et al. 2016). However, β-lyase activity varies considerably among brewing strains because of widespread inactivating mutations in the key β-lyase-encoding gene *IRC7* [15–17]. When such activities are seen in wine or beer fermentations, they are typically associated with wild strains [17, 18]. The β-lyase activity of *Saccharomyces* species outside of *S. cerevisiae* has not been explored much. From limited studies, it appears as if *Saccharomyces uvarum* strains tend to have higher thiol release abilities than *S. cerevisiae* strains [19, 20]. Interspecific hybridization between *S. cerevisiae* and *S. uvarum* strains has also been shown to enhance thiol release during wine fermentations in comparison to using *S. cerevisiae* by itself [20, 21]. A possible explanation for the enhanced β-lyase activity of *S. uvarum*, is that *IRC7* is located further away from the telomeres in *S. uvarum* compared to *S. cerevisiae*, and therefore less likely to be downregulated by sub-telomeric silencing as in *S. cerevisiae* [22].

*S. uvarum* is not traditionally used in beer fermentation, but is associated with wine and cider fermentations. It’s close ancestor, *Saccharomyces eubayanus*, is also not traditionally used in beer fermentations, however, the species has formed a interspecies hybrid with *S. cerevisiae* to give rise to lager yeast [23]. Recent studies have shown that both species do, however, have some brewing potential, as strains of both species tend to be able to utilize the main wort sugar maltose [24–27]. They are not capable of using all fermentable sugars though, and also possess other negative qualities from a brewing perspective, in particular the production of phenolic flavours. Studies have shown that these wild-yeast deficiencies can be overcome through yeast breeding [24, 28–30]. As mentioned above, breeding or hybridization has also been used to construct yeast strains with enhanced β-lyase activities [15, 31–33]. There is potential to create hybrid strains (*S. cerevisae* x *S. uvarum* or *S. cerevisae* x *S. eubayanus)* that incorporate the normal brewing yeast capabilities as well as an enhanced ability to release hop-derived flavours through enzymatic activity. Such strains could not only allow the introduction of new and enhanced flavour to beer, but also potentially yield cost and environmental impact savings from the need to use less hops to produce the same amount of flavour.

Here, we first screened the β-lyase activity of a set of potential parent strains. We then constructed interspecies hybrids between strains of *S. cerevisiae* and either *S. uvarum* or *S. eubayanus* using rare mating. Hybrids were then screened for fermentation ability and β-lyase activity. The hybrid with most efficient thiol release was phenolic off-flavour positive (POF^+^), so it was further mated with a POF^-^ *S. cerevisiae* parent strain to form a three-parent hybrid. This hybrid was sporulated and POF^-^ spore clones were successfully obtained. The fermentation performance and ability to release volatile thiols of this spore clone was demonstrated at 10L-scale. Our study demonstrates how the release of desirable hop-derived thiols in brewing yeast can be boosted by hybridization with wild yeast.

## Materials and methods

### Yeast strains

Lists of all yeast strains included in this study are found in Table 1 and 2. Table 1 contains a list of the included parent and control strains, while Table 2 contains a list of all constructed hybrid strains.

**Table 1.** A list of parent and control strains used in the study.

| Strain (short name) | Type | Description |
| --- | --- | --- |
| VTT A-75060 (A60) | *S. cerevisiae* | POF- maltotriose+ brewing strain |
| VTT A-81062 (A62) | *S. cerevisiae* | POF+ maltotriose+ brewing strain |
| VTT A-94132 (A132) | *S. cerevisiae* | POF- maltotriose- brewing strain |
| VTT C-12961 (C961) | *S. eubayanus* | POF+ maltotriose- wild strain |
| VTT C-201037 (C1037) | *S. uvarum* | POF+ maltotriose- wild strain |
| BFPBL32 (R-6.1) | *S. uvarum* | POF+ maltotriose- wild strain; Isolate in [67] |
| VTT A-63015 (A15) | *S. pastorianus* | Commercial lager yeast control |
| WLP001 | *S. cerevisiae* | Commercial ale yeast control; White Labs Inc. (USA) |

**Table 2.** A list of hybrid strains constructed in the study.

| Strain | Parent 1 | Parent 2 | Description |
| --- | --- | --- | --- |
| A60PA × C961L H5 | *S. cerevisiae* A60 | *S. eubayanus* C961 | Rare mating hybrid |
| A132U1 × C961 H7 | *S. cerevisiae* A132 | *S. eubayanus* C961 | Rare mating hybrid |
| A132U1 × C961L H9 | *S. cerevisiae* A132 | *S. eubayanus* C961 | Rare mating hybrid |
| A132U1 × C961L H10 | *S. cerevisiae* A132 | *S. eubayanus* C961 | Rare mating hybrid |
| A62U1 × C961L1 H11 | *S. cerevisiae* A62 | *S. eubayanus* C961 | Rare mating hybrid |
| A62U1 × C961L1 H12 | *S. cerevisiae* A62 | *S. eubayanus* C961 | Rare mating hybrid |
| A62U1 × C961L1 H13 | *S. cerevisiae* A62 | *S. eubayanus* C961 | Rare mating hybrid |
| A62U1 × C961L1 H14 | *S. cerevisiae* A62 | *S. eubayanus* C961 | Rare mating hybrid |
| A62U1 × C1037L1 L1 | *S. cerevisiae* A62 | *S. uvarum* C1037 | Rare mating hybrid |
| A62U1 × C1037L1 L2 | *S. cerevisiae* A62 | *S. uvarum* C1037 | Rare mating hybrid |
| A132s × Su32s C1 | *S. cerevisiae* A132 | *S. uvarum* BFPBL32 (R-6.1) | Spore-to-spore hybrid |
| A132s × Su32s D1 | *S. cerevisiae* A132 | *S. uvarum* BFPBL32 (R-6.1) | Spore-to-spore hybrid |
| A132 × (A62U1 × C1037L1 L2) D2 | *S. cerevisiae* A132 | Hybrid A62U1 × C1037L1 L2 | 'Three parent' hybrid |
| A132 × (A62U1 × C1037L1 L2) D2 C4 | *S. cerevisiae* A132 | Hybrid A62U1 × C1037L1 L2 | POF- spore clone of A132 × (A62U1 × C1037L1 L2) D2 |

### Hybrid construction

Interspecific hybrids between brewing and wild *Saccharomyces* strains were generated using both spore-to-spore mating and rare mating. Prior to rare mating, natural auxotrophic mutants (lys-or ura-) of the parental strains were selected on α-aminoadipic and 5-fluoroorotic acid agar plates, respectively [34, 35]. Auxotrophy was confirmed by the inability to grow on minimal selection agar medium (0.67 % Yeast Nitrogen Base without amino acids, 3 % glycerol, 3 % ethanol and 2 % agar).

For spore-to-spore mating, ascospores of the parental strains were generated by plating on 1% potassium acetate agar. Ascus walls were digested with 1 mg mL^−1^ Zymolyase 100T (US Biological, USA), after which spores from the different parental strains were dissected and placed next to each other on YPD agar plates (1 % yeast extract, 2 % peptone, 2 % glucose, and 2 % agar) using a micromanipulator (Singer Instruments, UK). The plates were incubated at 25 °C for 3 days, after which any emerging colonies were replated on minimal selection agar, and incubated at 25 °C for 5 days. Any colonies emerging on the minimal selection agar were regarded as potential hybrids

For rare mating, cultures of parent strains with complementary auxotrophy were grown overnight at 25 °C by inoculating a single colony into 50 mL of YPM (1 % yeast extract, 2 % peptone, 2 % maltose). One hundred microliters of the resulting overnight yeast cultures from both parental strains was transferred together to 1 mL YPM medium in a sterile 2 mL Eppendorf tube. Tubes were vortexed and incubated statically at 25 °C for 5 days. After incubation, the tubes were centrifuged at 5000×g for 5 min and the supernatant was removed. Five hundred microliters of starvation medium (0.1 % yeast extract and 0.1 % glucose) was added, and tubes were incubated for at least 2 h at room temperature. Tubes were then vortexed and 100 μL aliquots were spread onto minimal selection agar (without uracil or lysine). Plates were incubated at 25 °C for 5 days and any colonies emerging on the minimal selection agar were regarded as potential hybrids.

The three-parent hybrid D2 was constructed through spore-to-spore mating by using *S. cerevisiae* A132 and hybrid A62 × C1037 L2 as parent strains.

Selected hybrids were transferred to 1% potassium acetate agar for sporulation in an attempt to remove the POF phenotype. After 7 days of incubation at 25 °C, ascospores were digested using Zymolyase 100T (US Biological, Salem, MA, USA) and dissected on YPD agar using the MSM400 dissection microscope (Singer Instruments, Roadwater, UK).

### Hybrid confirmation

The hybrid status of potential hybrids was confirmed by PCR as described in Krogerus et al. (2015). Briefly, the rDNA ITS region was amplified using the primers ITS1 (5–-TCCGTAGGTGAACCTGCGG-3′) and ITS4 (5′-TCCTCCGCTTATTGATATGC-3′), and amplicons were digested using the *HaeIII* restriction enzyme (New England BioLabs, USA) as described previously [36]. The hybrid status of the potential three-parent hybrids was confirmed by PCR using delta12 (5′-TCAACAATGGAATCCCAAC-3′) and delta21 (5′-CATCTTAACACCGTATATGA-3′) primers.

### Enzyme activity of yeasts

β-lyase activity was estimated by measuring growth on cysteine. Media contained 0.17% Yeast Nitrogen Base without (NH_4_)_2_SO_4_ and amino acids, 1% glucose, 0.01% pyridoxal 5-phosphate, and 15 mM of cysteine. Growth assays were carried out in 96-well plates, with 145 μL media per well. Wells were inoculated (to a starting OD600 value of 0.1) with 5 μL of washed pre-culture suspended in water to an OD600 value of 3. Plates were sealed with a Breathe-Easy membrane (Sigma-Aldrich, Espoo, Finland), and incubated at 25 °C for one week. OD600 values were measured on a VarioSkan plate reader (Thermo Scientific, Vantaa, Finland), while cysteine content of the growth media was estimated using DTNB [37].

The ability to produce phenolic off-flavour was estimated using the absorbance-based method described by Mertens et al. (2017).

### Wort fermentations

Lab-scale wort fermentations were first carried out to screen the yeast hybrids in order to identify top-performing strains that were able to rapidly attenuate wort sugars. Fermentations were carried out in 100 mL Schott bottles capped with glycerol-filled airlocks. Yeast strains were grown overnight in 25 mL YPM medium at 25 °C. The pre-cultured yeast was then inoculated into 80 mL of all-malt wort (extract content of 15 °Plato) at a rate of 5 g fresh yeast L^−1^. Fermentations were carried out in duplicate at 25 °C for 7 days. Fermentations were monitored by mass lost as CO_2_.

2L-and 10L-scale wort fermentations were carried out in 3L and 12L cylindroconical stainless steel fermenting vessels, containing 2L of 15 °P wort or 10L of 12 °P wort. Yeast was propagated in autoclaved wort. The worts were produced at the VTT Pilot Brewery from barley malt and contained 2.5 g L^−1^ each of Cascade and Perle hops added to the whirlpool. The wort was oxygenated to 10 mg L^−1^ prior to pitching (Oxygen Indicator Model 26073 and Sensor 21158; Orbisphere Laboratories, Geneva, Switzerland). Yeast was inoculated at a rate of 10^6^ viable cells mL^−1^ °Plato^−1^, together with 5 g L^−1^ of Cascade hops (dry hopping). The fermentations were carried out in triplicate at 18 °C until no change in alcohol level was observed for 24 h or for a maximum of 9 days.

Wort samples were drawn regularly from the fermentation vessels aseptically and placed directly on ice, after which the yeast was separated from the fermenting wort by centrifugation (9000×g, 10 min, 1 °C).

### Beer chemical analysis

The specific gravity, alcohol level (% v/v), and pH of samples were determined from the centrifuged and degassed fermentation samples using an Anton Paar Density Metre DMA 5000 M with Alcolyzer Beer ME and pH ME modules (Anton Paar GmbH, Graz, Austria).

Concentrations of fermentable sugars (glucose, fructose, maltose, and maltotriose) and ethanol were measured by HPLC using a Waters 2695 Separation Module and Waters System Interphase Module liquid chromatograph coupled with a Waters 2414 differential refractometer (Waters Co., Milford, MA, USA). An Aminex HPX-87H Organic Acid Analysis Column (300 × 7.8 mm; Bio-Rad, Hercules, CA, USA) was equilibrated with 5 mM H_2_SO_4_ (Titrisol, Merck, Darmstadt, Germany) in water at 55 °C, and samples were eluted with 5 mM H_2_SO_4_ in water at a 0.3 mL min^−1^ flow rate.

Higher alcohols, esters and 4-vinyl guaiacol concentrations were analysed using headspace solid phase micro-extraction coupled with gas chromatography (Agilent 7890A)- mass spectrometry (Agilent 5975C; HS-SPME-GC-MS) by modifying the method used by Rodriguez-Bencomo et al. (2012). 4 mL of beer sample, 1.8 g of NaCl, and 50 μL of internal standard solution (containing 1.28 μg 3-octanol, 1.19 μg 3,4-dimethylphenol) were added to 20 mL headspace vials. The samples were pre-incubated in Gerstel MPS autosampler at 44.8 □ for 10 min and the volatiles were extracted by using a 2 cm divinylbenzene/ carboxen/polydimethylsiloxane (DVB/CAR/PDMS) fibre (Supelco) at 44.8 □ for 46.8 min. The samples were injected in splitless mode (10 min desorption time at 270 □) and the compounds were separated on an HP-Innowax silica capillary column (60 m, 0.250 mm i.d., 0.25 μm film thickness). The oven temperature program was from 40□°C (3□min) to 240□°C (4□°C□min^−1^) and the final temperature was held for 15 min. The MS data were collected at a mass range of 35-450 amu. Identification was based on spectral data of reference compounds and those of NIST 08 library. Calibration curves determined for 2-methoxy-4-vinylphenol, 2-phenylethyl acetate, 3-methylbutyl acetate, and ethyl esters of acetic, butyric, decanoic, hexanoic and octanoic acids, respectively (r^2^ = 0.933-0.999).

The volatile thiols 4-mercapto-4-methyl-2-pentanone (4MMP), 3-mercapto-1-hexanol (3MH), and 3-mercaptohexylacetate (3MHA) were determined at VLB Berlin using the method described by Dennenlöhr et al. (2020). In this method thiols are extracted and derivatized by headspace solid-phase microextraction (HS-SPME) with on-fiber derivatization (OFD) using 2,3,4,5,6-pentafluorobenzyl bromide (PFBBr). Resulting PFBBr-thioesters are then separated and analysed using gas chromatography tandem mass spectrometry (GC-MS/MS). The instrumental setup, parameters of sample preparation, GC-MS/MS analysis, calibration, and quantification were in accordance to Dennenlöhr et al. (2020), with an exception that the analysis was performed on an Agilent 7010B triple quadrupole MS with high-efficiency source instead of an Agilent 7000C triple quadrupole MS. Each sample was analysed in duplicate.

### Whole-genome sequencing and analysis

*S. cerevisiae* A132, *S. uvarum* C1037 and three hybrid strains were whole-genome sequenced inhouse. DNA was extracted using the method described by Denis et al. (2018). Sequencing was carried out on an Oxford Nanopore MinION MK1C instrument on a R10.3 flow cell using the SQK-LSK110 ligation sequencing kit. Reads were basecalled using Guppy version 5.0.11 using the ‘super high accuracy’ model. Basecalling statistics are available in Supplementary Table S1. Genomes of *S. cerevisiae* A132 and *S. uvarum* C1037 were *de novo* assembled using Flye version 2.9 [41]. Assemblies were polished using Medaka version 1.4.3. Reads from the hybrid strains were aligned to a concatenated reference genome of *S. cerevisiae* A81062 [42] and *S. uvarum* C1037 using minimap2 version 2.17-r941 [43]. Sequencing coverage was estimated with mosdepth (version 0.2.6; Pedersen and Quinlan 2018). Variants were called with clair3 [45]. Basecalled reads have been deposited in NCBI-SRA under BioProject number PRJNA820581.

## Results

### Screening β-lyase activity in parent strains

As the objective of the study was to construct brewing yeast strains with enhanced ability to release volatile thiols from wort, we first screened the β-lyase activity in a range of *Saccharomyces* strains. The set included *S. pastorianus* lager strains, *S. cerevisiae* ale strains, as well as *S. eubayanus* and *S. uvarum* wild strains. β-lyase activity was estimated based on the ability to grow on and consume 15 mM cysteine as the sole nitrogen source. This requires expression of a functional β-lyase-encoding *IRC7* gene in *S. cerevisiae*, and correlates with ability to release 4MMP from its cysteinylated form [15, 17, 46]. The strains showed variation in ability to grow and consume cysteine, with the *S. uvarum* strains showing the highest growth and consumption (Figure 1A and 1B). The poorest growth was observed for the lager and selected ale strain, while *S. cerevisiae* A60, *S. cerevisiae* A62 and *S. eubayanus* C961 showed intermediate growth and consumption. A strong positive correlation was observed between growth on 15 mM cysteine and its consumption (Figure 1C). Results suggest that hybridization of ale strains with either of the *S. uvarum* strains show highest potential for generating a hybrid strain with both efficient fermentation and ability to release volatile thiols.

**Figure 1.**
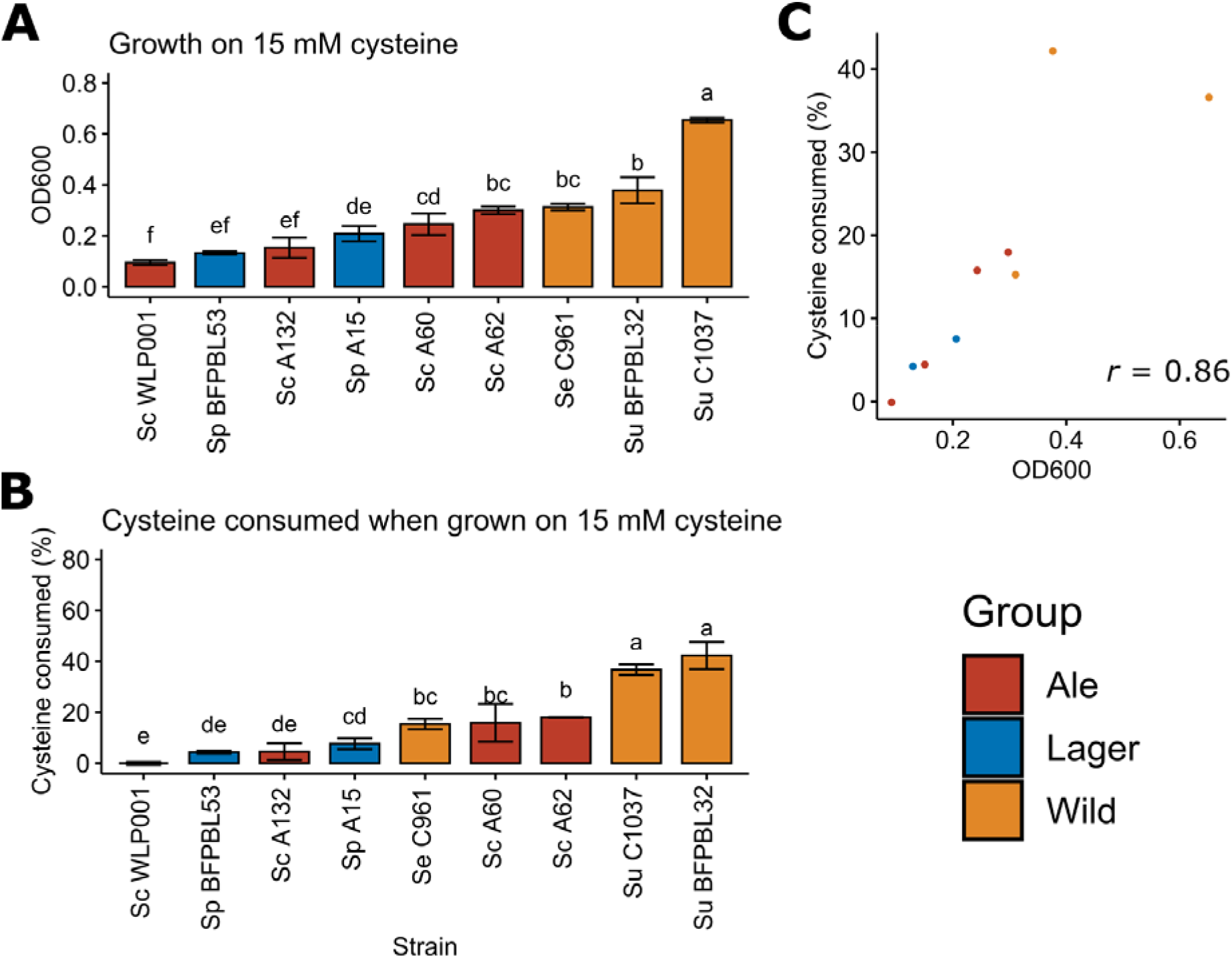
The ability to (**A**) grow on 15 mM cysteine as a sole nitrogen source and (**B**) consume cysteine (% of original amount) during cultivations. (**C**) The correlation and Pearson correlation coefficient (*r*) between growth and cysteine consumption. Strains are coloured based on type: blue, lager; red, ale; orange, wild. Different letters above bars indicate significant differences between values as determined by one-way ANOVA and Tukey’s post-hoc test (*p* < 0.05).

### Construction of interspecific hybrids

Following the initial screening, we attempted to construct interspecies hybrids between one of three *S. cerevisiae* ale strains (A60, A62 and A132) and one of three wild *Saccharomyces* strains (*S. uvarum* BFPBL32, *S. uvarum* C1037 and *S. eubayanus* C961). Hybrid construction was first attempted with rare mating, to ensure the whole genome of both parent strains were to be retained. As rare mating requires selection markers for the parent strains, we selected auxotrophic or petite mutants of the parent strains prior to mating. Uracil auxotrophs of *S. cerevisiae* A62 and A132, a petite mutant of *S. cerevisiae* A60, as well as lysine auxotrophs of *S. eubayanus* C961 and *S. uvarum* C1037 were successfully obtained. No auxotrophs of *S. uvarum* BFPBL32 were obtained, and the strain had to be excluded from the rare mating attempts. Rare mating resulted in ten successful hybrids of four different hybrid combinations (see Table 1). Hybrid status was confirmed using PCR, showing presence of genetic material from both parent strains (Figure 2).

**Figure 2.**
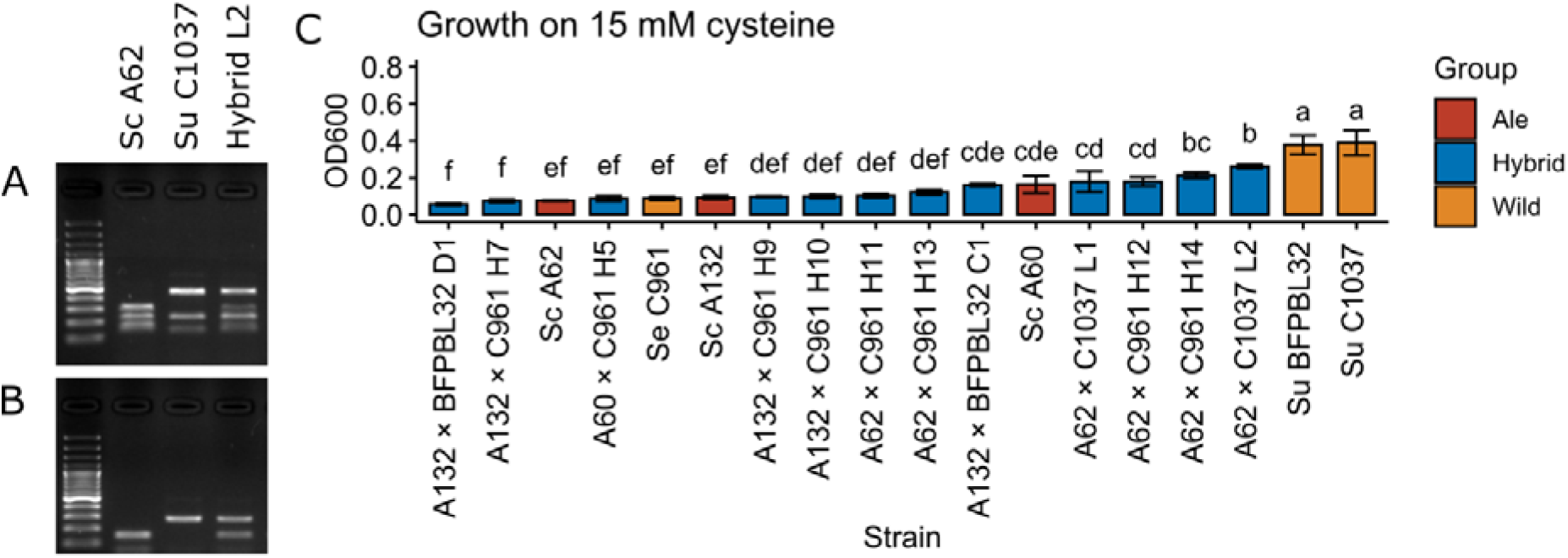
Confirmation of hybridization by (**A**) rDNA ITS PCR and RFLP, and (**B**) amplification of DBP6 and MEX67 genes. Lane 1 is a 100 bp DNA ladder, lane 2 *S. cerevisiae* A62, lane 3 *S. uvarum* C1037 and lane 4 an interspecies hybrid between the two. (**C**) The ability to grow on 15 mM cysteine as a sole nitrogen source. Strains are coloured based on type: blue, hybrid; red, ale; orange, wild. Different letters above bars indicate significant differences between values as determined by one-way ANOVA and Tukey’s post-hoc test (*p* < 0.05).

As not all hybrid combination were successful, it was further attempted to construct hybrids using spore-to-spore mating. This technique does not require selection of auxotrophic mutants of the parent strains prior to mating, but the parent strains need to be able to produce viable spores. Of the three brewing strains, only *S. cerevisiae* A132 sporulated efficiently. *S. uvarum* C1037 also did not sporulate. As hybrids between A132 and C961 were already obtained with rare mating, spore-to-spore mating was ultimately only tested between *S. cerevisiae* A132 and *S. uvarum* BFPBL32. Two successful hybrids were obtained (see Table 1).

### Hybrid screening

Following successful construction of interspecific hybrids, the twelve hybrid strains were screened for β-lyase activity in the same way as the parent strains. Variation between the hybrids was observed, and strains tended to exhibit mid-parent heterosis (values in between parent strains) in regards to ability to grow on 15 mM cysteine as a sole nitrogen source (Figure 2C). The highest growth among the hybrids was observed for the A62 × C1037 L2.

After screening β-lyase activity, 80 mL-scale 15 °Plato wort fermentations were carried out with 8 hybrids and the 6 parent strains to identify hybrids with good fermentation performance. Most hybrids had fermentation rates in between those of the parent strains, but certain hybrids outperformed both parent strains (Figure 3A-C). These include the A60 × C961 hybrid and several hybrids created with *S. cerevisiae* A62. Similar results, of hybrids outperforming the parent strains in regards to fermentation, have been observed before in multiple hybrid combinations [15, 24, 28, 47]. However, on average, there was no significant difference between the final attenuation reached by the parents or hybrids (Figure 3D). Two hybrids were selected for 2L-scale fermentations based on fermentation performance and β-lyase activity.

**Figure 3.**
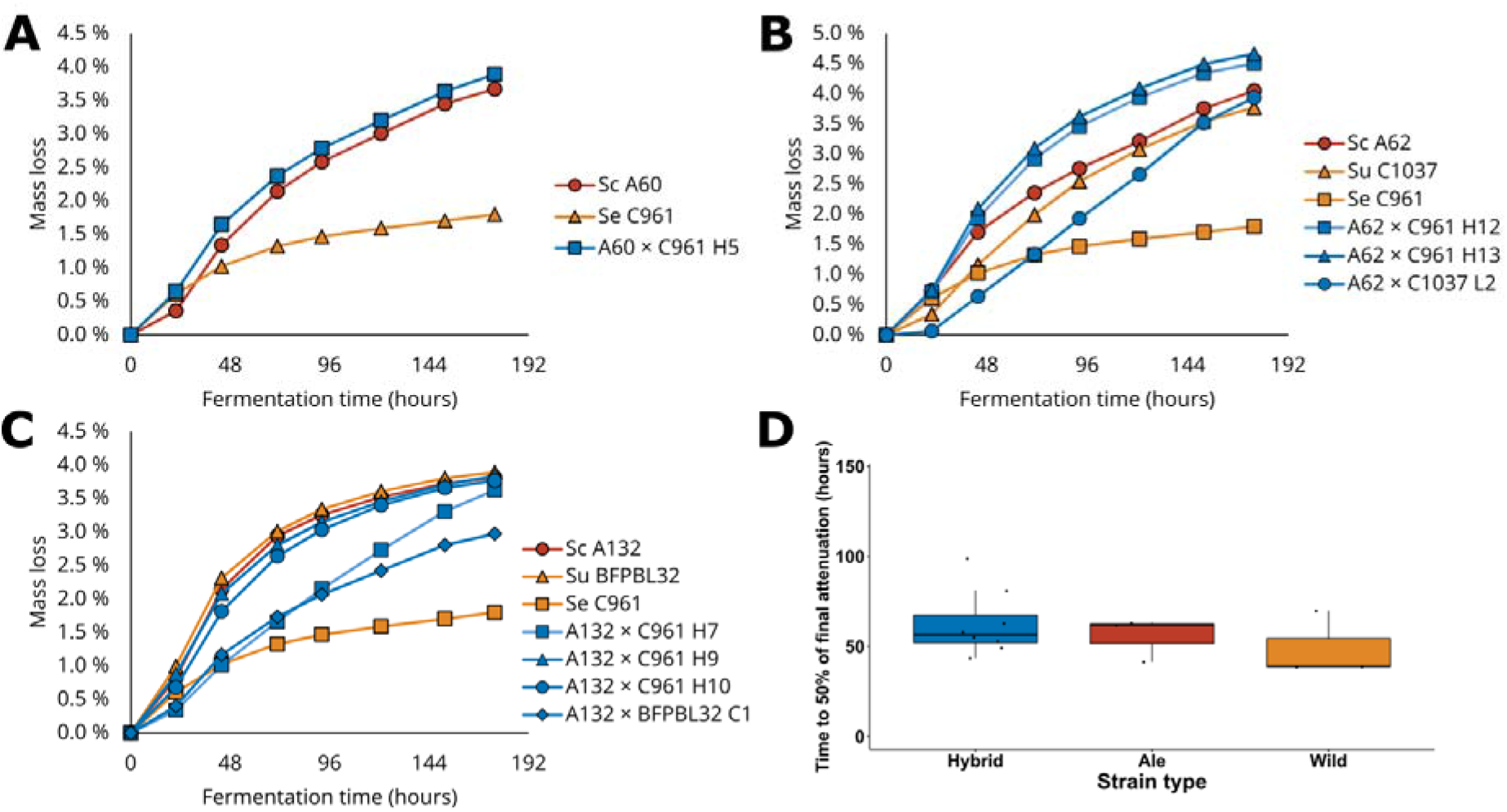
The fermentation performance (% mass lost as CO_2_) of *de novo* hybrids compared **to** parent strains during 80 mL-scale fermentation with 15 °Plato wort. (**A-C**) Strains are plotted **in** separate graphs based on the *S. cerevisiae* brewing parent: (**A**) *S. cerevisiae* A60, (**B**) **S.** cerevisiae A62, and (**C**) *S. cerevisiae* A132. (**D**) The average time (hours) until 50% of final attenuation was reached in the three strain groups. Strains are coloured based on type: blue, hybrid; red, ale; orange, wild.

### Wort fermentations confirm enhanced phenotypes in hybrids

2L-scale fermentations in 15 °Plato wort were performed with the two selected hybrids (A62 × C1037 L2 and A132 × BFPBL32 C1) and the four parent strains that were used for their creation. Fermentation performance at 2 L-scale was similar to that during the smaller scale fermentations (Figure 4A and B). One hybrid (A62 × C1037 L2) fermented faster than both parent strains, while the other fermented slower than both parent strains (A132 × BFPBL32 C1). Of the hybrids, only A62 × C1037 L2 was able to consume maltotriose. This trait was inherited from *S. cerevisiae* A62, which was also the only parent strain capable of maltotriose use.

**Figure 4.**
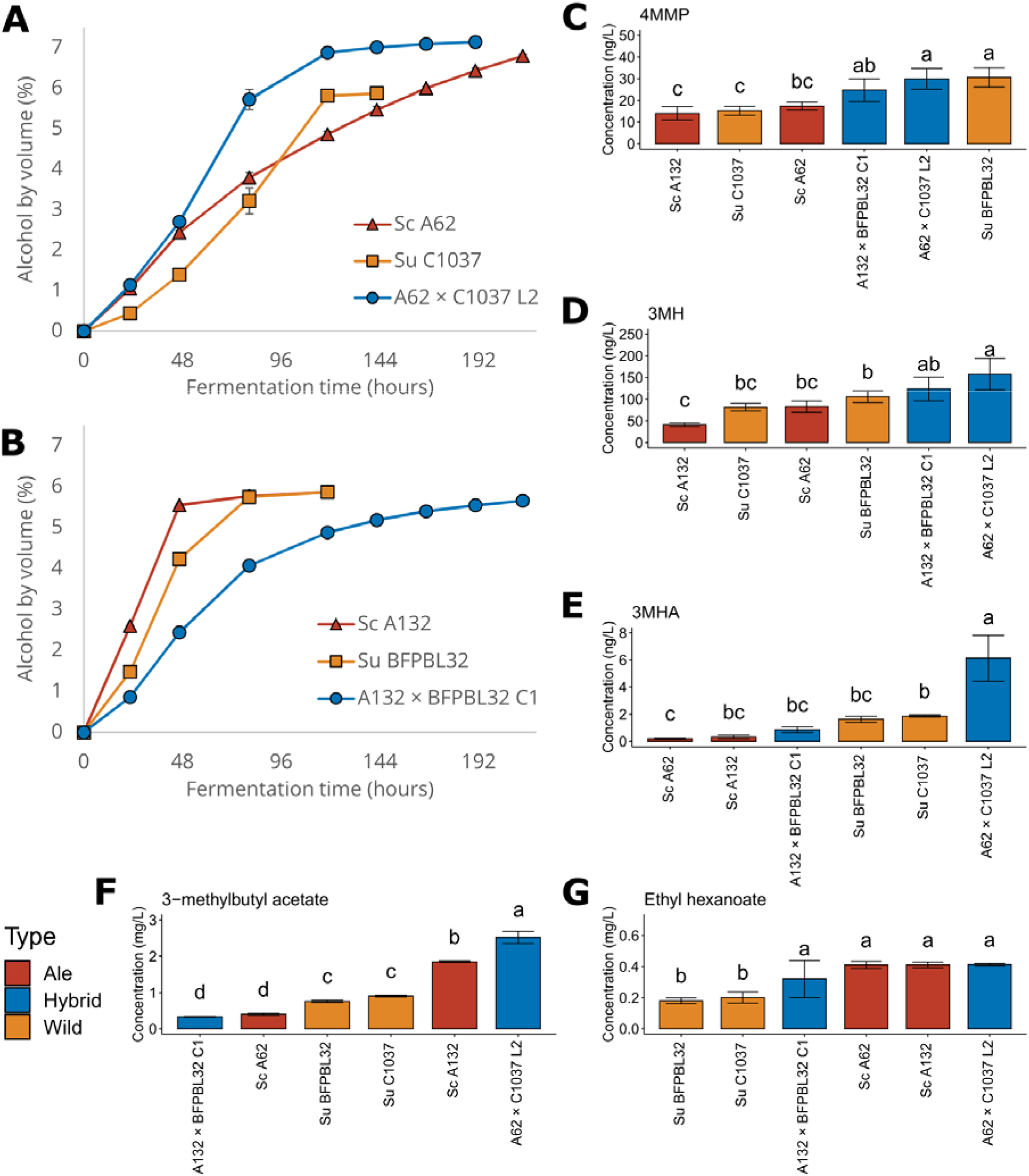
The fermentation performance of *de novo* hybrids compared to parent strains during **2** L-scale fermentation with 15 °Plato wort. (**A-B**) The alcohol by volume (%) produced durin**g** fermentation by (**A**) Hybrid A62 × C1037 L2 and (**B**) Hybrid A132 × BFPBL32 C1 compared to their respective parent strains. The concentrations (ng/L) of (**C**) 4-mercapto-4-methyl-2-pentanon**e** (4MMP), (**D**) 3-mercapto-1-hexanol (3MH), and (**E**) 3-mercaptohexyl acetate (3MHA) in th**e** resulting beers. The concentrations (mg/L) of (**F**) 3-methylbutyl acetate and (**G**) ethyl hexanoate in the resulting beers. Curves and bars are coloured based on type: blue, hybrid; red, ale; orang**e,** wild. Different letters above bars indicate significant differences between values as determined **by** one-way ANOVA and Tukey’s post-hoc test (*p* < 0.05).

The beers showed significant variation in concentrations of volatile thiols (Figure 4C-E). Interestingly, the hybrids tended to show either higher or as high concentrations of 4MMP, 3MH and 3MH-acetate compared to the better parent (i.e. the wild *Saccharomyces* strain). Thiol levels had increased many fold from those measured in the wort, indicating β-lyase activity during fermentation (data not shown). Concentrations of 4MMP and 3MH were in almost all cases above the flavour threshold of 1 ng/L and 60 ng/L, respectively [14]. The A62 × C1037 hybrid, which had exhibited the most efficient fermentation, also produced beer with the highest levels of volatile thiols. Hence, the objective of creating a strain with both efficient fermentation and an enhanced ability to release hop-derived thiols was successfully reached.

The beers also varied in their aroma-active ester content (Figure 4F and G). The A62 × C1037 hybrid again exhibited best-parent heterosis and produced higher levels of esters compared to either parent. For the A132 × BFPBL32 hybrid, concentrations of 3-methylbutyl acetate and ethyl hexanoate were lower than and in between the parents, respectively. The beer made with the A62 × C1037 hybrid had both the highest levels of volatile thiols and aroma-active esters, and therefore appeared to produce the most fruity and complex aroma. However, as both parent strains of this hybrid were POF^+^, the hybrid was also POF^+^. This trait is undesired in most beer styles, and we thus next attempted to remove the phenotype.

### Construction of three-parent hybrid and POF^−^ spore clone

To further develop the most promising, but POF^+^, hybrid A62 × C1037, an attempt was made to mate the hybrid with a POF^−^ *S. cerevisiae* A132 parent strain. This was accomplished through spore-to-spore mating. As the hybrid was tetraploid (being a result of rare mating), it was able to sporulate and produce viable spores despite being an interspecies hybrid. These spores were crossed with spores of *S. cerevisiae* A132 to form a three-parent hybrid. Hybrid status was assessed using interdelta PCR, and a single confirmed hybrid, A132 × (A62 × C1037 L2) D2, was obtained (Supplementary Figure S1).

As the three-parent hybrid was still POF^+^, from containing functional *PAD1* and *FDC1* genes from the A62 × C1037 hybrid, it was also sporulated in an attempt to obtain POF^−^ spore clones. Spore viability was low, and only 4 out of 24 possible spore clones were isolated. These spore clones were screened for the POF phenotype, and three out of four spore clones were POF^−^ (Figure 5B). Two of the three POF^−^ spore clones also showed the highest apparent β-lyase activity as well (Figure 5A).

**Figure 5.**
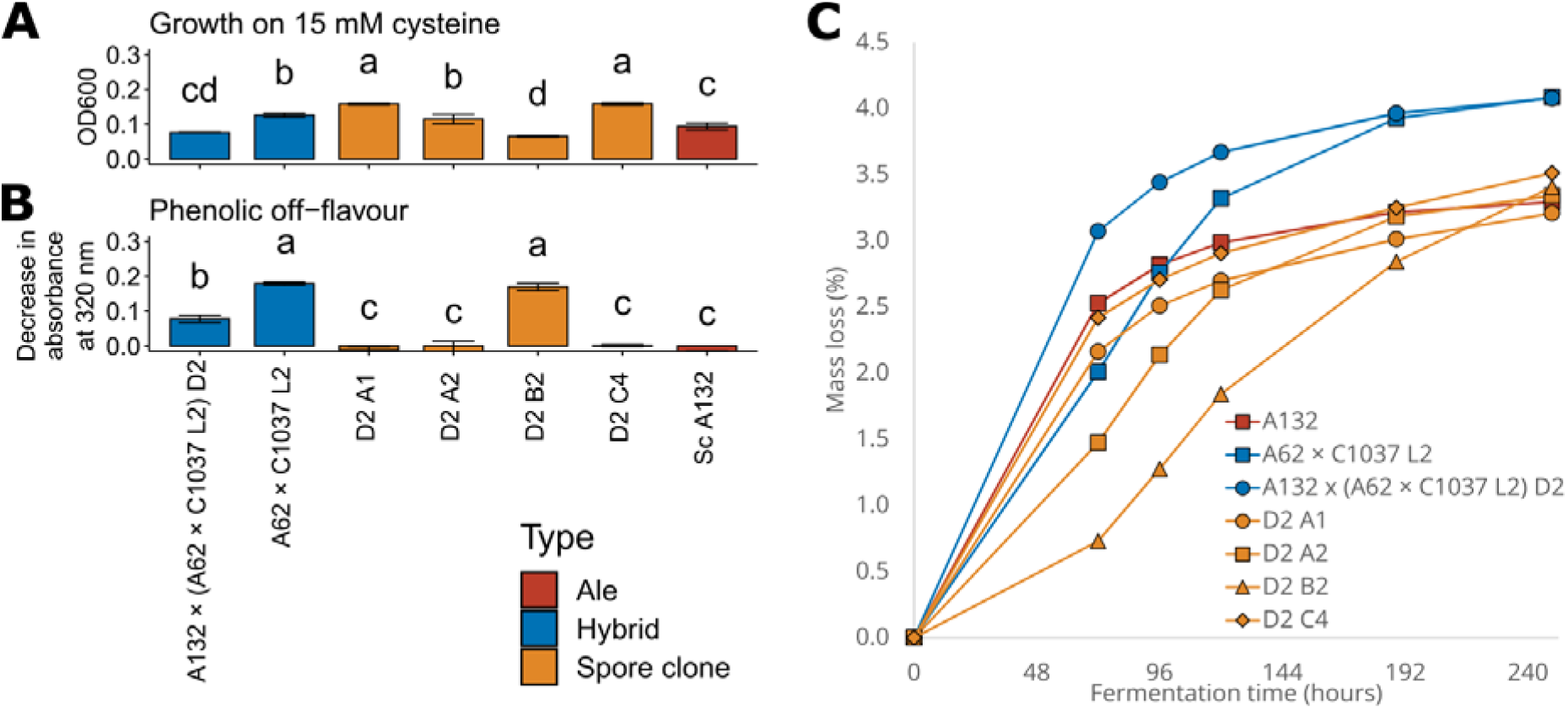
The ability of the three-parent hybrid and derived spore clones to (**A**) grow on 15 mM cysteine as a sole nitrogen source and (**B**) produce phenolic off-flavour by converting ferulic ac**id** (decrease in absorbance at 320 nm). (**C**) The mass lost as CO_2_ (%) during 80 mL-scale fermentation of 15 °Plato wort with parent strain, hybrids, and four spore clones. Strains are coloured based on type: blue, lager; red, ale; orange, spore clone. Different letters above bars indicate significant differences between values as determined by one-way ANOVA and Tukey’s post-hoc test (*p* < 0.05).

Small-scale wort fermentations were also performed to ensure sufficient fermentation performance of the spore clones. The three-parent hybrid fermented the fastest and reached the highest attenuation, followed by the original A62 × C1037 hybrid (Figure 5C). The spore clones fermented the slowest out of the compared strains, but ‘D2 C4’ was identified as the spore clone with the best fermentation performance. This strain was also POF^−^ and showed good β-lyase activity. The strain fermented as fast as the *S. cerevisiae* A132 parent strain, but reached a higher attenuation thanks to the ability to use maltotriose.

Whole-genome sequencing of hybrids A62 × C1037 L2, A132 × (A62 × C1037 L2) D2, and A132 × (A62 × C1037 L2) D2 C4 was also performed to examine the genome structure of these hybrids and confirm mutations in the POF-related genes. *De novo* assemblies of the parent strains *S. cerevisiae* A132 and *S. uvarum* C1037 were first generated from long sequencing reads, while a *de novo* assembly of *S. cerevisiae* A62 was obtained from Krogerus et al. (2021b). The resulting assemblies of 48 and 19 scaffolds for *S. cerevisiae* A132 and *S. uvarum* C1037 (including the 16 chromosomes and mitochondrial DNA) and spanned genome sizes of 12.5 and 12.1 Mbp, respectively (assembly statistics and dot plots are available in Supplementary Table S2 and Supplementary Figures S2-3). For analysis of the hybrid strains, a concatenated reference genome of *S. cerevisiae* A62 and *S. uvarum* C1037 was used. The A62 × C1037 L2 hybrid produced through rare mating appeared to be tetraploid with nearly two copies of all chromosomes derived from *S. cerevisiae* and *S. uvarum* (Figure 6A). The three-parent hybrid A132 × (A62 × C1037 L2) D2 also appeared to be tetraploid, but rather had approximately three copies of all *S. cerevisiae* chromosomes and one copy of *S. uvarum* chromosomes (Figure 6B). This is expected from spore to spore mating of the tetraploid *S. cerevisiae* A132 and the tetraploid A62 × C1037 L2 hybrid. Finally, the POF^−^ spore clone A132 × (A62 × C1037 L2) D2 C4 appeared diploid, and the strain had lost numerous *S. uvarum* chromosomes completely, while most *S. cerevisiae* chromosomes were present in two copies (Figure 6C). This included *S. uvarum* chromosome 12, which carries the *S. uvarum* orthologue of *IRC7*, and *S. uvarum* chromosome 13, which carries the *S. uvarum* orthologues of *PAD1* and *FDC1*. The *S. cerevisiae* A132 parent strain carries a loss-of-function and frameshift mutation (460C>T, Gln154* and 501dupA, Trp168fs, respectively) in the *FDC1* gene, rendering the strain POF^−^. The same mutations were found in the three-parent hybrid A132 × (A62 × C1037 L2) D2 and its POF^−^ spore clone D2 C4 (Figure 6D). However, they were heterozygous in D2 (i.e. functional alleles of *FDC1* were still present), while homozygous in D2 C4 (i.e. only non-functional present). In both the *S. cerevisiae* parent strains A62 and A132, the β-lyase coding *IRC7* gene carries an inactivating 38-bp deletion. However, the deletion is heterozygous in A62, meaning the full-length *IRC7* was present in one copy in all three hybrid strains L2, D2 and D2 C4 (Figure 6E).

**Figure 6.**
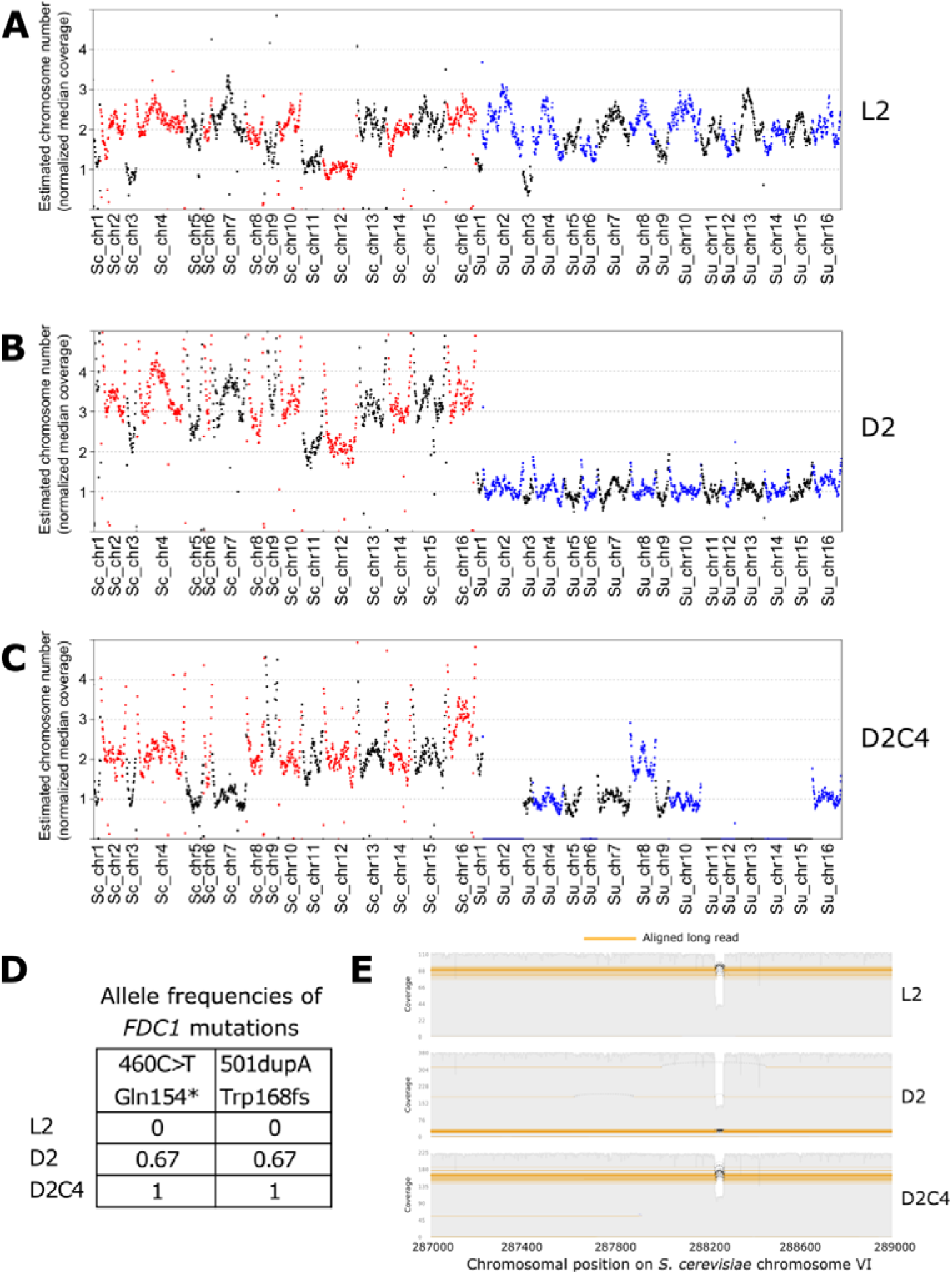
Whole genome sequencing of three hybrid strains. (**A**-**C**) The estimated chromosome copy numbers of hybrid strains hybrids (**A**) A62 × C1037 L2, (**B**) A132 × (A62 × C1037 L2) D2, and (**C**) A132 × (A62 × C1037 L2) D2 C4. The copy numbers were estimated by normalizing th**e** median sequencing coverage in 10 kb windows to the median sequencing coverage across the whole genome. *S. cerevisiae* and *S. uvarum* chromosomes are coloured alternating red-black and blue-black, respectively. (**D**) The allele frequencies of nonsense mutations in *S. cerevisiae FDC1* detected in the three hybrid strains. (**E**) The sequencing coverage along *S. cerevisiae IRC7* in the three hybrid strains. The inactivating heterozygous 38-bp deletion occurs around position 288200 along chromosome VI.

### 10 L-scale fermentations confirm enhanced phenotypes in hybrid and spore clone

To confirm the fermentation phenotypes in larger-scale fermentations, we performed 10L-scale fermentations in 12 °Plato wort with one parent strain (*S. cerevisiae* A62) along with two hybrids (A62 × C1037 and the POF- ‘D2 C4’ spore clone). Fermentations proceeded as observed previously at smaller scale, with all strains ultimately reaching the same attenuation, but the hybrids reaching it slightly faster (Figure 7A). The high attenuation (>85 %) indicated maltotriose use by all strains, and this was confirmed by HPLC analysis (data not shown).

**Figure 7.**
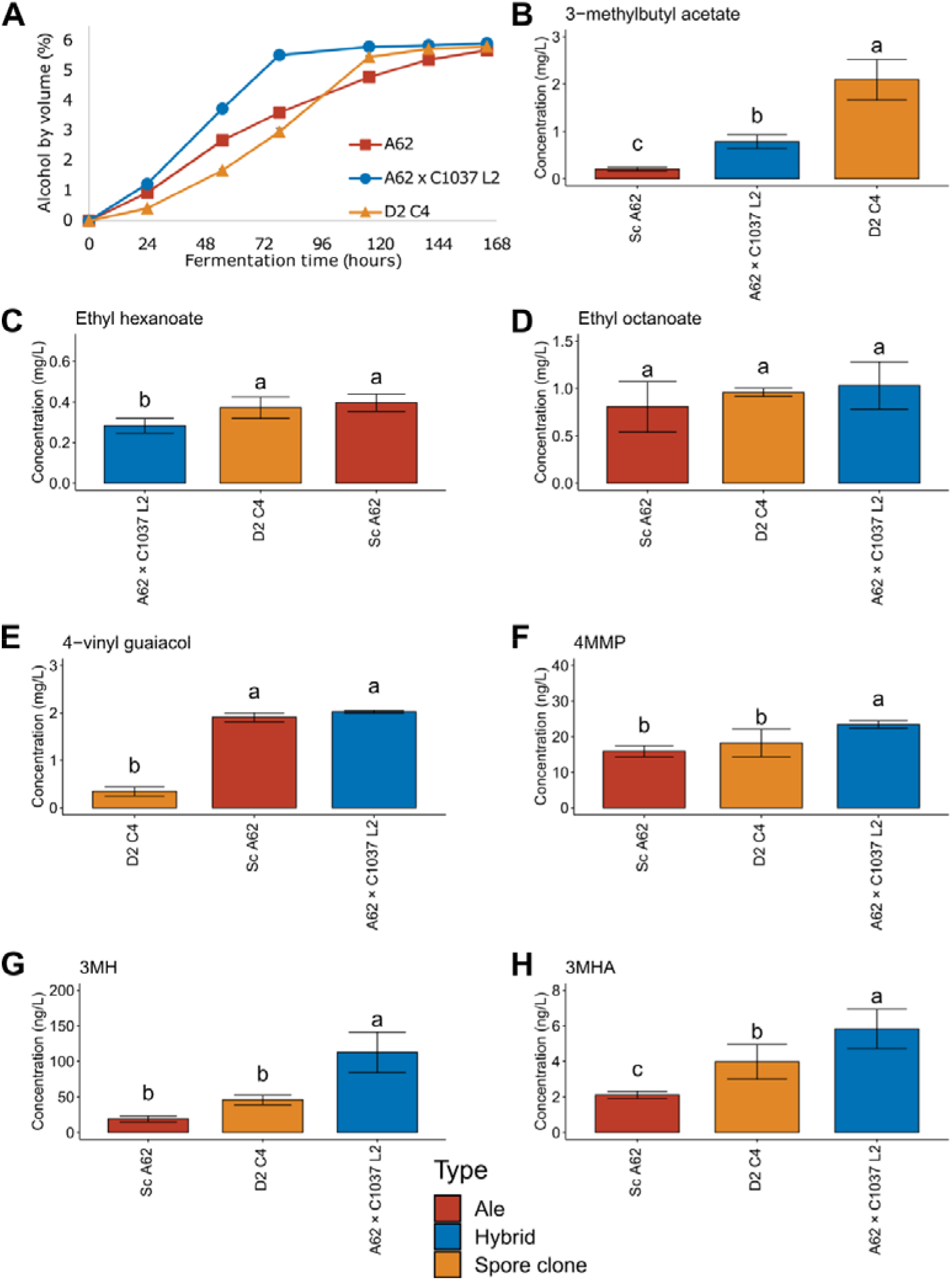
Fermentation performance and concentrations of thiols, esters and 4-vinylguaicol in 10 L-scale fermentations of 12 °Plato wort. (**A**) Alcohol by volume (%) during fermentation**s.** Concentrations (mg/L) of (**B**) 3-methylbutyl acetate, (**C**) ethyl hexanoate, (**D**) ethyl octanoate, an**d** (**E**) 4-vinyl guaiacol in the beers. Concentrations (ng/L) of (**F**) 4-mercapto-4-methyl-2-pentanon**e** (4MMP), (**G**) 3-mercaptohexanol (3MH), and (**H**) 3-mercaptohexylacetate (3MHA) in the beers. Different letters indicate significant differences (*p* < 0.05) as determined by one-way ANOVA and Tukey’s post-hoc test.

In regards to aroma formation, significantly less 4-vinyl guaiacol was found in the beer produced with the ‘D2 C4’ spore clone (Figure 7E). Hence, confirming the POF phenotype that was observed earlier during the POF assay and genome sequencing. In regards to aroma-active esters, the beers made with the hybrid strains contained significantly higher concentrations of 3-methylbutyl acetate, while few differences were observed for the ethyl esters (Figure 7B-D). The ‘D2 C4’ spore clone was therefore not only POF^−^, but also produced more desirable fruity esters than the other strains here.

Like with the esters, thiols levels were also higher in the beers produced with the hybrid strains (Figure 7F-H). The beers fermented with the hybrid strains showed significantly higher concentrations of 4MMP, 3MH and 3MHA compared to the parent strain (no difference in 4MMP concentration between A62 and ‘D2 C4’). In the beer fermented with the hybrid strains, concentrations of all three thiols were above or around their flavour thresholds of 1, 60, and 4 ng/L for 4MMP, 3MH, and 3MHA, respectively [14]. This suggested that the thiols had a positive influence on flavour. Thiol levels were lower in the beer fermented with the spore clone, compared to that with the A62 × C1037 hybrid. As β-lyase activity was strongly linked with the *S. uvarum* parent, it is likely that this decrease in thiol release is due to the loss of *S. uvarum IRC7*.

## Discussion

As a result of growing demand for highly hopped beer with fruity aroma, much attention has been placed on maximising and modifying the aroma obtained from hops. In addition to adjusting hop amount, hop variety and timing of the hop additions, the aroma compounds derived from the hops can also be modified through yeast-mediated enzymatic reactions. One such class of compounds are volatile thiols, which have been shown to play a central role in contributing fruity hop aroma to beer [12, 48–50]. While these thiols can transfer directly from the hops to beer, the vast majority of the volatile thiols present in hops are found in odorless cysteinylated or glutathionylated forms [11, 51, 52]. However, free thiols can be liberated from these conjugated precursors through the action of β-lyase enzymes [13, 53, 54], thus releasing higher concentrations of thiols into the beer from the same amount of hops.

Here, we aimed to generate brewing yeast strains with enhanced ability to release volatile thiols during wort fermentation through interspecific hybridization of *S. cerevisiae* brewing strains with strains of *S. uvarum* or *S. eubayanus*. Interspecific hybridization has been used extensively for brewing yeast strain development in the past decade [28–30, 55, 56]. It allows for combining traits from diverse parent strains, can be performed without any genetic engineering, and hybrids may in some cases exhibit hybrid vigour, where they outperform the parent strains in certain traits [7]. The approach has been used, for example, to construct interspecific brewing hybrids with both enhanced wort fermentation and a more diverse aroma profile [24, 29]. The hybrids generated in this study were able to release higher concentrations of 4MMP and 3MH than their *S. cerevisiae* parent, and in some cases also the *S. uvarum* parent. The result is in line with a study comparing the performance of a range of *S. cerevisiae* × *S. uvarum* hybrids in wine fermentations [33]. There, they observed higher release of 4MMP among the *S. uvarum* and hybrid strains compared to the *S.cerevisiae* strains. In addition, we also recently observed enhanced thiol release during beer fermentations with selected intraspecific *S. cerevisiae* hybrids generated between brewing and wild strains of *S. cerevisiae* [15]. Interspecific hybridization can therefore be used to enhance thiol release during beer fermentations as well.

In *Saccharomyces* yeasts, β-lyase enzymes responsible for releasing 4MMP and 3MH are coded by *IRC7* and *STR3* genes [13, 53, 54, 57]. *IRC7*, in particular, has been shown to be key for thiol release, and the gene is well studied in *S. cerevisiae* [13, 17, 31, 58]. Irc7p activity varies greatly between strains, and a number of inactivating mutations have been identified [13, 17]. These are particularly prevalent among domesticated strains, and include a 38-bp deletion that significantly decreases enzyme activity [17, 18]. Here, we observed the presence of the 38-bp inactivating deletion in both *S. cerevisiae* A132 (homozygous) and A62 (heterozygous), which explains the lower thiol concentrations in the beers fermented with these strains. It is unclear what causes the higher β-lyase activity of the *S. uvarum* strains that was observed here, but it could be simply related to a full-length allele of *IRC7*, or that the *S. uvarum* orthologue encodes a protein with higher activity. The activity of *S. uvarum* β-lyase enzymes have, to our knowledge, not been studied or compared to *S. cerevisiae* orthologues. Another possibility is that the enhanced β-lyase activity of *S. uvarum* is related to *IRC7* regulation and its genomic position [22]. In *S. cerevisiae, IRC7* is located in the sub-telomeric region of chromosome VI (around 5 kbp from the right telomere), and this region is downregulated through silencing by the *SIR2*-encoded histone deacetylase [59, 60]. In our *de novo* assembly of *S. uvarum* C1037, *IRC7* was located in the sub-telomeric region of chromosome XII (around 18 kbp from the left end of the assembled chromosome). It remains to be explored in future studies how these locations affect *IRC7* expression.

Thiol release during beer fermentations is also affected by process conditions. In a recent study with lager yeast, it was revealed that fermentation temperature has a contrasting effect on the release of cysteinylated compared to glutathionylated thiols [61]. Release of thiols from glutathionylated precursors was enhanced at higher fermentations (24 °C), while release from cysteinylated precursors was higher at lower fermentations (12 °C). Lager yeasts, i.e. natural *S. cerevisiae* × *S. eubayanus* hybrids, are known for their cold tolerance, and fermentations are usually carried out at lower temperatures than those with *S. cerevisiae* ale yeast [62]. This cold tolerance is inherited from the *S. eubayanus* parent [25, 28]. *S. uvarum* strains typically also exhibit enhanced cold tolerance compared to *S. cerevisiae* [24, 63, 64]. While not tested here, interspecific hybrids tend to have a broader temperature range for growth and fermentation compared to the parent strains alone [29, 30, 47, 65]. This could be exploited by using the constructed hybrids for both low and high temperature wort fermentations, in order to release different profiles of thiols. Using a low-temperature maturation phase (4 °C) after the primary fermentation was also recently revealed to increase the release of thiols into the beer [66], and *S. eubayanus, S. uvarum* and their hybrids are more likely to be metabolically active at such low temperatures compared to *S. cerevisiae* ale strains.

To conclude, our study reveals that interspecific hybridization can be used to enhance the β-lyase activity and thiol releasing ability of brewing yeast strains. We observed higher β-lyase activity in the *non-cerevisiae Saccharomyces* strains compared to *S. cerevisiae* brewing strains, and hybrids between them were shown to release high amounts of volatile thiols during fermentation. While the study here only focused on interspecific hybrids with *S. uvarum* and *S. eubayanus*, it is possible that other species in the *Saccharomyces* genus also exhibit higher β-lyase activity compared to *S. cerevisiae* brewing yeast strains. This could be explored in future studies. As has been seen in previous studies, the constructed hybrid strains here also showed improved fermentation and formation of yeast-derived aroma compounds. Unwanted traits, such as phenolic off-flavour production, could be further removed by exploiting the fertility of the allotetraploid hybrids. Increased thiol release has many advantages for the fermentation of modern hoppy beer. It not only allows for the production of beer with boosted hop aroma, but decreases the environmental impact of the brewing process through lower hop use.

## Acknowledgements

We thank Aila Siltala, Niklas Fred, Eero Mattila and Ronja Eerikäinen for technical assistance. Benjamin Palmer and Sarah Thörner are acknowledged for thiol analysis.

## Declarations

### Availability of data and material

The ONT reads generated in this study have been submitted to NCBI-SRA under BioProject number PRJNA820581 in the NCBI BioProject database (https://www.ncbi.nlm.nih.gov/bioproject/).

### Authors’ contributions

KK: Conceived the study, designed experiments, performed experiments, analysed all data, wrote the manuscript.

NR: Supervised thiol analysis, edited the manuscript.

BG: Conceived the study, designed experiments, edited the manuscript.

All authors read and approved the final manuscript.

### Ethics approval

This article does not contain any studies with human participants or animals performed by any of the authors.

