## Supplementary Material for "Increased volatile thiol release during beer fermentation using constructed interspecies yeast hybrids"

**Table S1.** Statistics of basecalled long sequencing reads.

|  | **A132** | **C1037** | **L2** | **D2** | **D2C4** |
| --- | --- | --- | --- | --- | --- |
| Mean read length | 44322.9 | 37418.9 | 43705.7 | 37018.5 | 31709.4 |
| Mean read quality | 13.9 | 14.1 | 13.3 | 13.2 | 13.5 |
| Median read length | 25463 | 18244 | 18496 | 21503.5 | 18156.5 |
| Median read quality | 14.1 | 14.3 | 13.4 | 13.3 | 13.6 |
| Number of reads | 27162 | 41754 | 51878 | 106910 | 78564 |
| Read length N50 | 83510 | 77444 | 98218 | 66148 | 56753 |
| Total bases | 1203899231 | 1562387095 | 2267362140 | 3957648463 | 2491214502 |

**Table S2.** Statistics of *de novo* assemblies of *S. cerevisiae* A132 and *S. uvarum* C1037.

|  | **A132** | **C1037** |
| --- | --- | --- |
| Total sequence count | 48 | 19 |
| Total sequence length | 12516089 | 12124555 |
| Min sequence length | 556 | 14033 |
| Max sequence length | 1514186 | 1297703 |
| Mean sequence length | 260751.9 | 638134.5 |
| Median sequence length | 25599 | 650706 |
| N50 | 929913 | 917883 |
| L50 | 6 | 6 |
| N90 | 267256 | 480116 |
| L90 | 15 | 13 |
| A% | 30.92 | 29.97 |
| T% | 30.88 | 29.91 |
| G% | 18.93 | 19.97 |
| C% | 19.03 | 20.11 |
| AT% | 61.8 | 59.87 |
| GC% | 37.96 | 40.09 |
| N% | 0.24 | 0.04 |

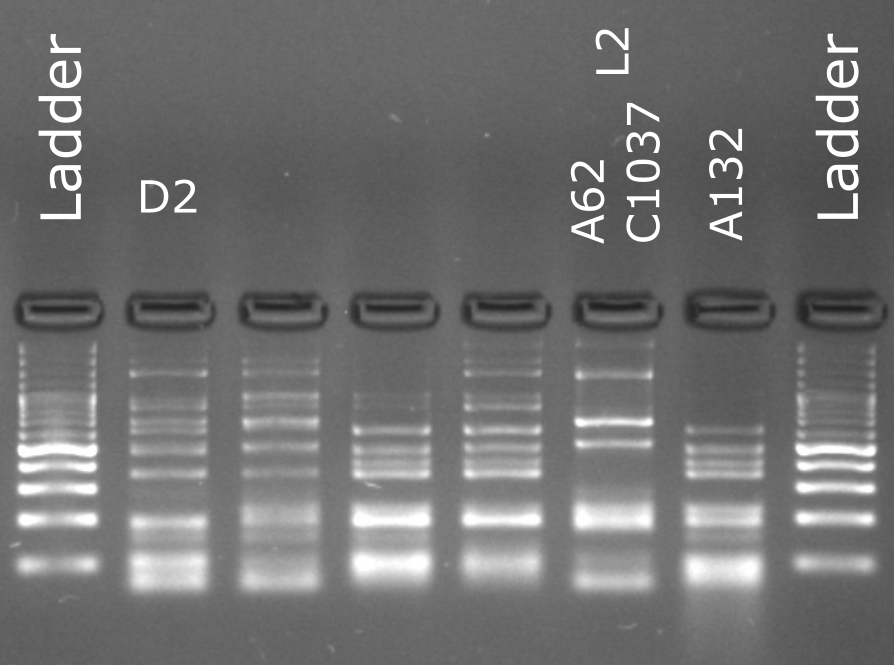

**Figure S1** - Confirmation of successful hybridization of A132 × (A62U1 × C1037L1 L2) D2 (lane labelled ‘D2’) from parent strains A62U1 × C1037L1 L2 (lane labelled ‘A62 C1037 L2’) and *S. cerevisiae* A132 (lane labelled ‘A132’) using interdelta PCR. Hybridization was also later confirmed separately with whole genome sequencing.

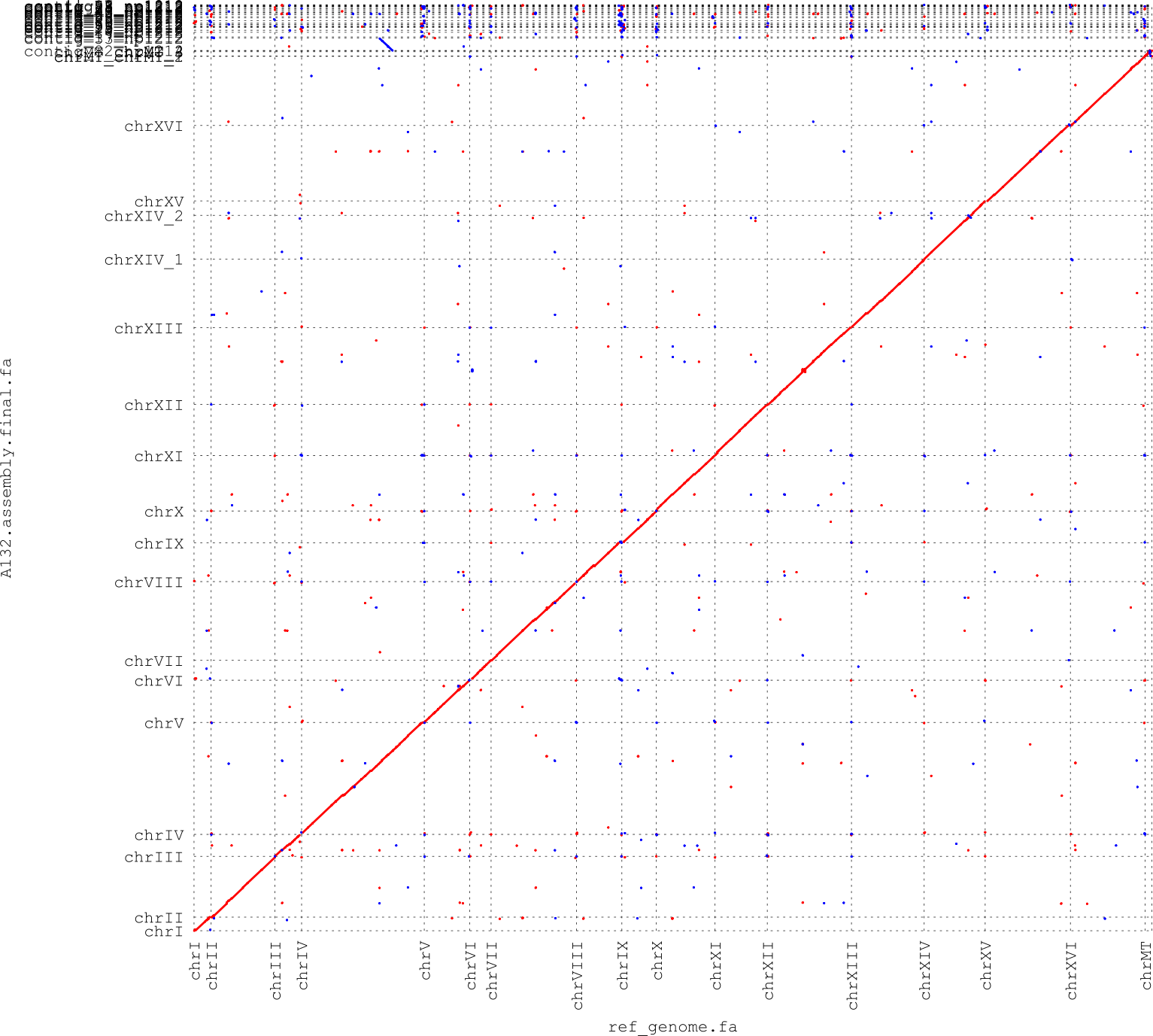

**Figure S2** - Alignment of *S. cerevisiae* A132 assembly contigs (y-axis) to *S. cerevisiae* S288C (x-axis)

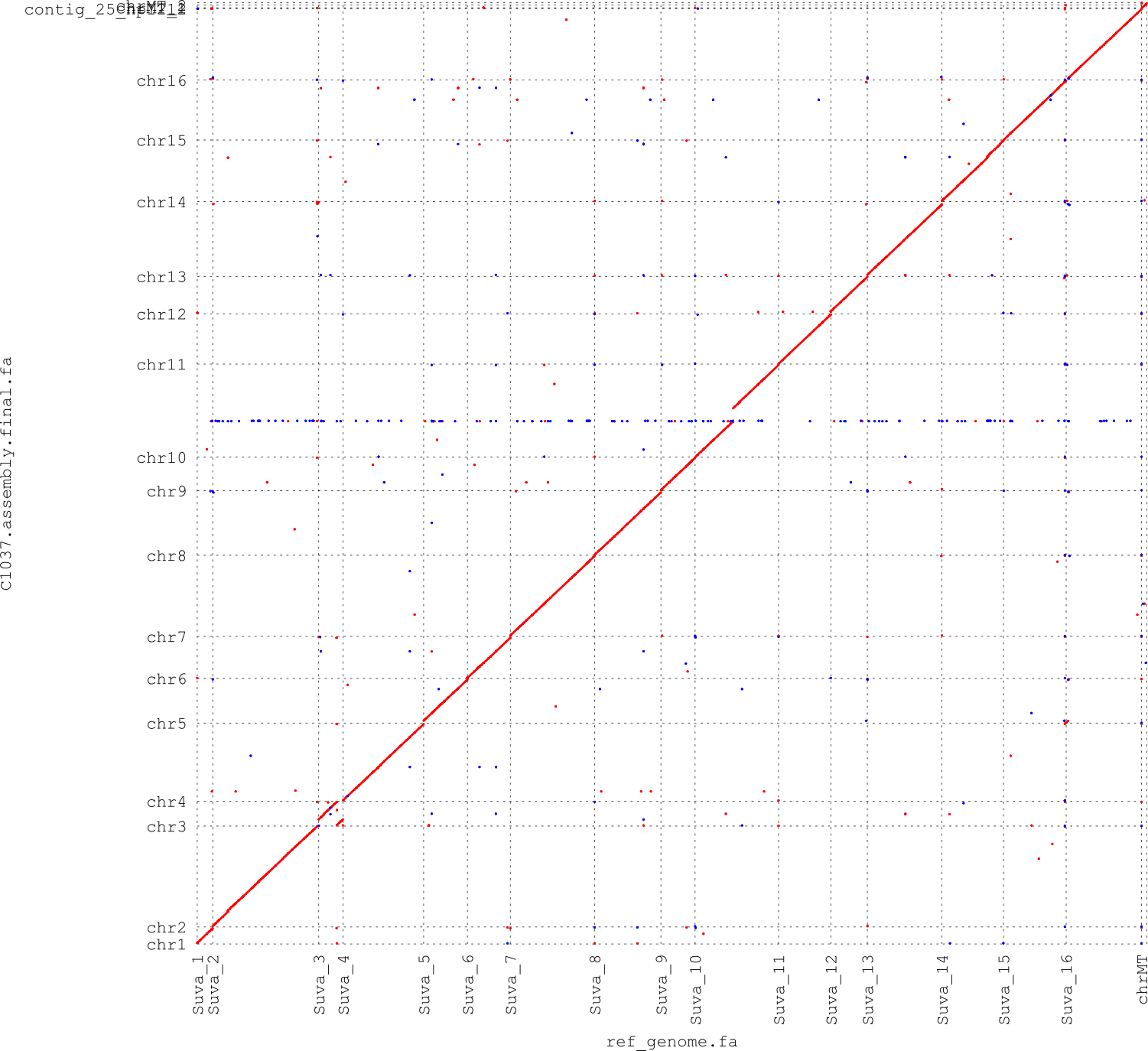

**Figure S3** - Alignment of *S. uvarum* C1037 assembly contigs (y-axis) to *S. uvarum* CBS 7001 (x-axis)
